# Usefulness of a familiarity signal during recognition depends on test format: Neurocognitive evidence for a core assumption of the CLS framework

**DOI:** 10.1101/797837

**Authors:** Regine Bader, Axel Mecklinger, Patric Meyer

## Abstract

Familiarity-based discrimination between studied target items and similar foils in yes/no recognition memory tests is relatively poor. According to the complementary learning systems (CLS) framework this is due do a relatively small difference in familiarity strength between these two item classes. The model, however, also predicts that when targets and corresponding similar foils are presented next to each other in a forced-choice corresponding (FCC) test format, familiarity values for targets and foils can be directly compared because in each trial, targets are reliably more familiar than their corresponding foils. In contrast, when forced-choice displays contain non-corresponding foils (FCNC) which are similar to other studied items (but not the target), familiarity should not be diagnostic because familiarity values are not directly comparable (as in yes/no-tasks). We compared ERP old/new effects (ERPs of targets vs. foils) when participants were tested with FCC vs. FCNC displays after having intentionally encoded pictures of objects. As predicted, the mid-frontal old/new effect which is associated with familiarity was significantly larger in FCC compared to FCNC displays. Moreover, the target-foil amplitude difference predicted the accuracy of the recognition judgment in a given trial. This is one of the very few studies which support the assumption of the CLS framework that the test format can influence the diagnosticity of familiarity. Moreover, it implies that the mid-frontal old/new effect does not reflect the mean difference in the familiarity signal itself between studied and non-studied items but reflects the task-adequate assessment of the familiarity signal.

## Introduction

It is well established that recognition memory is generally supported by two distinct processes. While familiarity is a mere feeling of having something encountered before, recollection involves memory for details of a prior encounter (Yonelinas, 2002). Although the capability to recognize events on the basis of familiarity is generally impressive, it breaks down when studied and non-studied items are too similar (e.g., Morcom, 2015). A computational explanation for this is given by the complementary learning systems (CLS) framework (Norman & O’Reilly, 2003): Recollection is assumed to rely on the integrity of the hippocampus, which assigns pattern-separated (i.e. non-overlapping) representations to each single episode, even when events are similar. In contrast, familiarity signals are assumed to be created by the medial temporal lobe cortex (MTLC), which assigns overlapping representations to similar events. Thus, although studied items are more familiar than non-studied similar foils, the difference in familiarity strength between these two item classes is on average relatively small and their familiarity strength distributions are highly overlapping (see Figure 1) rendering familiarity-based recognition unreliable (Migo, Montaldi, & Mayes, 2009). This usually leads to high error rates in standard yes/no (YN) tasks, where test items are presented one at a time and a global decision criterion for all test trials can be assumed. However, the CLS predicts better performance when studied targets and corresponding similar foils are presented together on a forced-choice (FC) test display because their familiarity strength values can be directly compared permitting trial-unique decision criteria.

**Figure 1.**
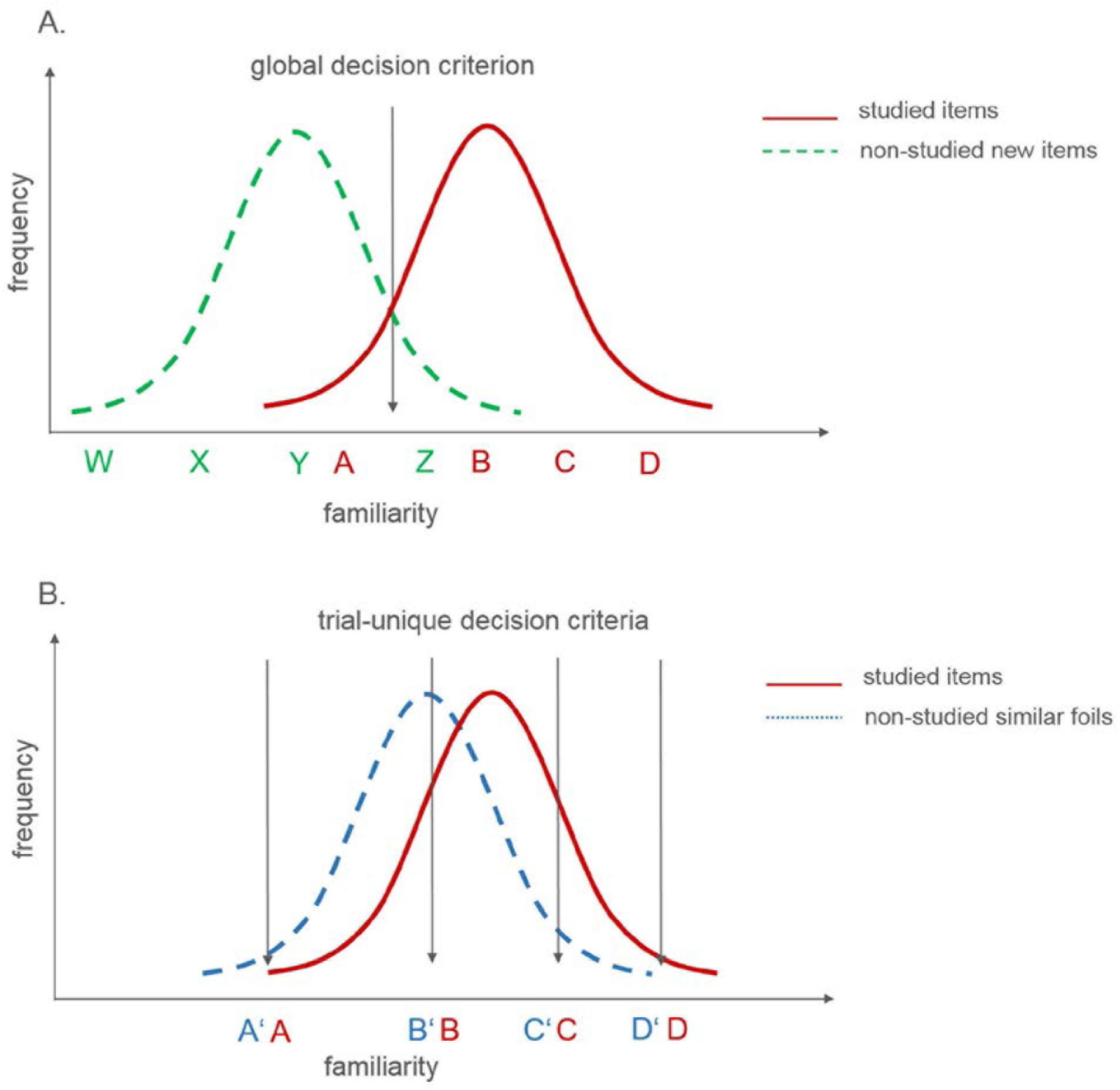
Familiarity distributions of studied and non-studied items during test assuming equal variances. A. When studied items (A,B,C,D) and non-studied new items (W,X,Y,Z) are dissimilar, familiarity distributions are only partly overlapping and a global decision criterion (as assumed in yes-no-tests) can be used. B. When studied items (A,B,C,D) and non-studied foils (A’,B’,C’,D’) are similar, familiarity differences due to study exposure are smaller than overall variance leading to strongly overlapping distributions and thus only the use of trial-specific decision criteria as in forced-choice corresponding tests is useful.

In line with this CLS assumption, patient Y.R., who had a selective hippocampal lesion, which impaired recollection but spared familiarity, performed within the range of healthy controls in a picture recognition test involving similar foils when tested in a FC but not in a YN test (Holdstock et al., 2002). While other studies showed no benefit from FC tests for hippocampal patients (Bayley, Wixted, Hopkins, & Squire, 2008; Jeneson, Kirwan, Hopkins, Wixted, & Squire, 2010), one study with older adults, for whom a disproportional deficit in recollection is assumed, showed an increase in familiarity-based responses in a FC compared to a YN test when recognition memory for similar faces was tested (Bastin & van der Linden, 2003). Moreover, a recent study (Migo et al., 2009) that investigated test format effects on familiarity-based recognition in healthy younger individuals contrasted three conditions: YN, forced-choice with targets next to corresponding similar foils (FCC), and forced-choice with targets next to foils which were similar to other studied items (forced-choice non-corresponding, FCNC). The FCNC condition serves as control condition because it is comparable with FCC tests (Figure 2A), but does not allow direct comparison of familiarity strength values. Supporting the view that familiarity is more useful in the FCC condition, instructions to use only familiarity reduced performance in FCNC and YN conditions compared to standard instructions, but not in the FCC condition.

**Figure 2.**
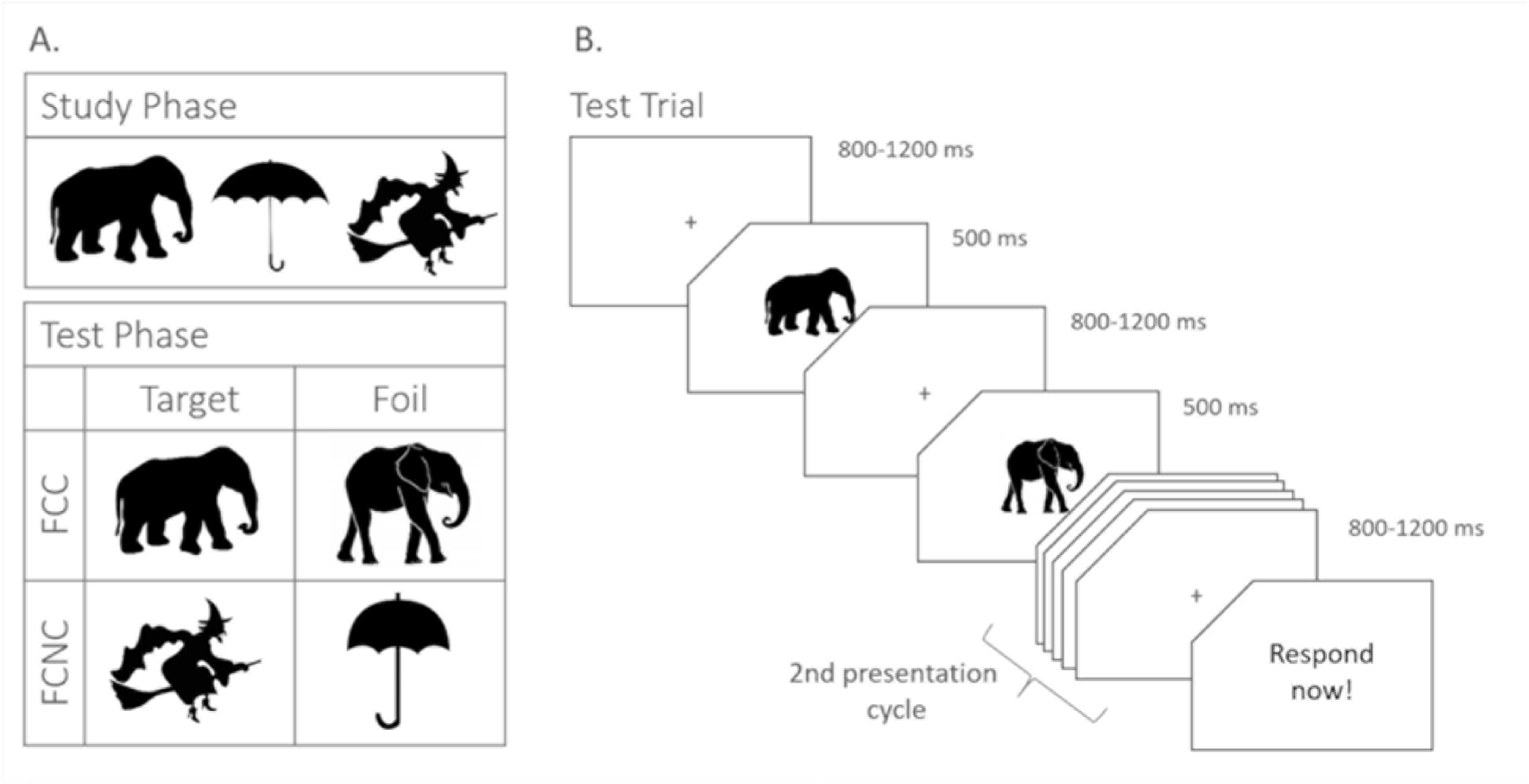
Experimental materials and trial procedure. A. Example target-foil combinations in forced-choice corresponding (FCC) and forced-choice non-corresponding (FCNC) conditions. B. Schematic illustration of a test trial in the FCC condition. Presentation was identical in the FCNC condition except for the target-foil combination. The first four displays of the trial were repeated in the 2^nd^ presentation cycle. Targets were at the 1^st^ position in half of the trials for each participant.

As neuropsychological and behavioral studies revealed mixed results, neurocognitive evidence in healthy young participants is essential, but still missing. Here, we explored the effects of test format on familiarity and recollection using event-related potentials (ERPs). Typically, ERPs to old items are more positive-going than those to new items. The mid-frontal old/new effect (300-500 ms post-stimulus) has been associated with familiarity while recollection has been linked to the later occurring (500-800 ms) left-parietal old/new effect (Rugg & Curran, 2007; but see Paller, Voss, & Boehm, 2007). FCC and FCNC conditions were administered in two study-test-cycles. The intentional study phases included a shoebox task for black and white pictures. As ERPs in the test phase were recorded separately for targets and foils, they had to be presented sequentially. In standard FC procedures, participants presumably switch back and forth between the two simultaneously presented pictures. Thus, in order to enhance the comparability of the sequential with the simultaneous presentation format and to also enable back and forth switching between pictures, the target-foil sequence was repeated within a trial allowing participants to look at each picture twice (Figure 2B; see Voss & Paller, 2009, for a similar procedure). In addition, differences between conditions become evident not before both pictures have been presented at least once as only the second of each picture pair reveals whether the two pictures are similar or not. Hence, our analyses focused on the second presentation of both pictures. In line with the CLS, we predicted that the mid-frontal old/new effect would be larger in the FCC than FCNC condition. In a second step, we used a logistic regression approach to assess whether the amplitude difference between the target and the foil for each single trial reflects the diagnostic value of the familiarity signal and thus can predict the accuracy of a subject’s response (see Noh, Liao, Mollison, Curran, & De Sa, 2018; Ratcliff, Sederberg, Smith, & Childers, 2016, for other studies using single trial approaches).

## Materials & Methods

### Participants

Thirty-two students from Saarland University (16 female, mean age of 24.4 (19-35) years) participated in the experiment. Two additional subjects had to be excluded because they did not perform above chance (p(hits) > .5) in the recognition test as revealed by a binomial test (p > .05) or not enough artefact-free trials (at least 13 per condition). All participants were right-handed as assessed by the Edinburgh Handedness Inventory (Oldfield, 1971), had normal or corrected-to-normal vision and no known neurological problems (self-report). They gave informed consent and were reimbursed with eight Euros/hour or course credit. The study was approved by the local ethics committee and adhered to the Declaration of Helsinki.

### Stimuli & Procedure

Visual stimuli were 352 pairs of black and white images (silhouettes or icons) collected from the internet. Each pair consisted of two very similar versions of one object. The experiment was divided into two study-test blocks, one for each condition. The order of the blocks was counterbalanced. In each study phase, participants were presented with 176 single images. In the FCNC condition, 88 of these images were presented as targets in the test phase together with a foil which was similar to one of the remaining 88 images form the study phase. Thus, in the FCNC condition, only one version of each pair appeared during the test as judgments on one image of a similar pair could influence subsequent judgments on the other one. In the FCC condition, 88 images were presented in the test phase together with the corresponding similar foil. The remaining 88 images from the study phase served as filler items to equalize block length in both conditions because in the FCNC condition 176 study pairs were needed to obtain 88 test trials (see Figure 2A).

During the study phase, participants had the task to memorize the images and to decide whether the depicted object was smaller or larger than a shoebox. A study trial started with a 500 ms fixation cross before the image which was presented for 3000 ms. Afterwards a question mark prompted participants to make the shoebox decision for which they had a maximum of 1500 ms. After a 500 ms blank screen the next trial started. After every 22 trials participants could make a self-paced short break.

Each test trial comprised two presentation sequences of target and foil. To avoid EEG oscillations time-locked to stimulus presentation, each sequence started with a jittered fixation cross (800-1200 ms) followed by the presentation of the first image for 500 ms. After another jittered fixation cross (800-1200 ms), the second image was shown for 500 ms. For each participant, the target was the first image within this sequence for half of the trials and the second image for the other half. After the second presentation of the sequence, a jittered fixation cross (800-1200 ms) appeared followed by a prompt (“Jetzt antworten!” / “Respond now!”) to indicate whether the target was the first or the second image within the sequence. Participants had a maximum of 1000 ms to respond. After a 1000 ms blank screen the next trial began. After every 22 trials participants could make a self-paced break.

Before the first block, there was a short study-test practice block to familiarize the participants with the complete procedure and to make the two blocks as equal as possible. In the FCNC block, subjects were explicitly told that foils were similar to previously studied images.

### EEG Data Acquisition & Processing

BrainVision Recorder 1.0 (Brain Products) was used to record EEG continuously from 59 scalp sites according to the extended 10-20 system (Jasper, 1958). The EEG was amplified with electrode AFz as ground electrode and the left mastoid electrode as reference using a 16-bit BrainAmp Amplifier (Brain Products). Data were digitized using a sampling rate of 500 Hz and an on-line analog band-pass filter of .016 to 250 Hz. Data were stored using an on-line digital low-pass filter of 100 Hz. Impedances were kept below 5 kΩ. Four additional electrodes were placed on the outer canthi and above and below the right eye to record electrooculographic (EOG) activity. BrainVision Analyzer 2.1 (Brain Products) was used for offline data processing which started with visually discarding excessive artefacts to improve independent component analysis (ICA) performance which was used for the correction of EOG and cardiac artifacts. The EEG was first filtered using a .05-30 Hz Butterworth filter (order: 4) and the ICA with a classic restricted infomax algorithm was employed. After re-referencing to the average of both mastoid electrodes, the data were segmented into −200 to 1000 ms epochs relative to image onset for each of the four image presentations within a trial. The epochs were baseline-corrected and artifacts were rejected by identifying segments including voltage steps greater than 30 μV/ms, voltage differences greater than 100 μV within a 200 ms interval or greater absolute amplitudes than +/- 70 μV. Finally, the data were checked manually for remaining artifacts (especially excessive alpha waves). For graphical illustration, filtered (12 Hz low-pass) waveforms were exported and we used the ggplot2 package of the software R (Wickham, 2009) to plot the ERP waveforms. Brain Vision Analyzer 2.1. (Brain Products) was used to create topographic maps.

### Experimental Design and Statistical Analysis

Inferential statistics were conducted using the software R (R Core Team, 2017) in RStudio (RStudio Team, 2016), especially the packages tidyverse (Wickham, 2017), lme4 (Bates, Maechler, Bolker, & Walker, 2015), lmerTest (Kuznetsova, Brockhoff, & Christensen, 2017) and ez (Lawrence, 2016). Significance level was set to α = .05. Behavioral data were analyzed with paired t-test comparing the two conditions (FCC, FCNC). For the ERP data, only data from the second presentation cycle were entered in the analyses. Based on previous literature (e.g., Küper, Groh-Bordin, Zimmer, & Ecker, 2012; Morcom, 2015), mean amplitudes from 300-500 ms were extracted for the mid-frontal old/new effect and for the sake of completeness from 500-650 ms for the left-parietal old/new effect. The latter time window was slightly shorter compared to other ERP recognition memory studies because of the offset potential being evident in the ERP around 700 ms. Amplitudes were pooled to be analyzed in a fronto-central (F1, Fz, F2, FC1, FCz, FC2) and a centro-parietal cluster (CP1, CPz, CP2, P1, Pz, P2). To examine the effect of test format on the old/new effects, only ERPs from correctly answered trials were used. Mean trial numbers per condition ranged from 33 to 37. As outlined in the Introduction, analyses were restricted to the second presentation cycle. The 2 x 2 x 2 repeated-measures ANOVAs included the factors Condition (FCC, FCNC), Item Type (target, foil), and Picture Position (first, second). Only effects involving the factors of Condition or Item Type are reported. Significant interactions were followed-up with t-tests for which in case of unplanned comparisons p-values were adjusted according to the Bonferroni-Holm procedure (Holm, 1979). In case of directional hypotheses, we report the p-values of the one-tailed test. Partial eta square 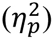 and Cohen’s d_av_ with the average of the two standard deviations as the denominator are provided as measures of effect size.

In order to test the hypothesis that accuracy of a participant’s response in a single trial can be predicted based on the ERP signature of familiarity strength in a single trial, we used multi-level binary logistic regression instead of standard binary logistic regression analyses to account for dependencies in the data within subjects. For each trial, difference scores between target and foil amplitudes in the fronto-central ROI were calculated for the early time window (see Rosburg, Mecklinger, & Frings, 2011, for a similar approach). These difference scores were entered into two different multi-level models per condition. For one model, we included the target-foil difference scores as a predictor and permitted random intercepts across subjects (random intercepts only model). In the other model, we allowed also the predictor to vary across subjects (random intercepts / random slopes model). We then compared whether the random intercepts / random slopes model reliably improved the fit of the data as compared to the random intercepts only model based on a chi-square test of the change in −2 log likelihood. In case of no improvement, we kept the simpler model. Significance of single effects was assessed based on the significance test for the predictor (z-statistic). To test whether the target-foil difference score better predicted the accuracy of a response in the FCC than the FCNC condition, we used the data of both conditions and included the difference score, condition and the interaction term of condition x difference score in the model. Before this, we centered the difference score variable within subjects and used centered values for condition (−1 = FCC, 1 = FCNC).

## Results

### Behavioral Results

Paired t-tests revealed that overall accuracy was significantly higher in the FCC condition (*M* = .82, *SD* = .09) than in the FCNC condition (*M* = .75, *SD* = .13; t(31) = 4.34, p < .001, d_av_ = 0.63) whereas mean reaction times were only marginally significantly smaller in the FCC condition (*M* = 323, *SD* = 69) than in the FCNC condition (*M* = 341, *SD* = 78; t(31) = 1.79, p = .083, d_av_ = 0.24).

### ERP Results

As depicted in Figure 3, which shows ERP waveforms collapsed across picture positions, ERPs in the early time window at frontal electrodes are generally more positive in the FCC condition than in the FCNC condition. Moreover, targets elicit more positive-going waveforms than foil items in the FCC condition while no such difference is observable for the FCNC condition. As can be seen in Figure 4.B, this is especially evident during the first picture position of the second cycle. A similar pattern in the waveforms is observable for the late time window and parietal recording sites.

**Figure 3.**
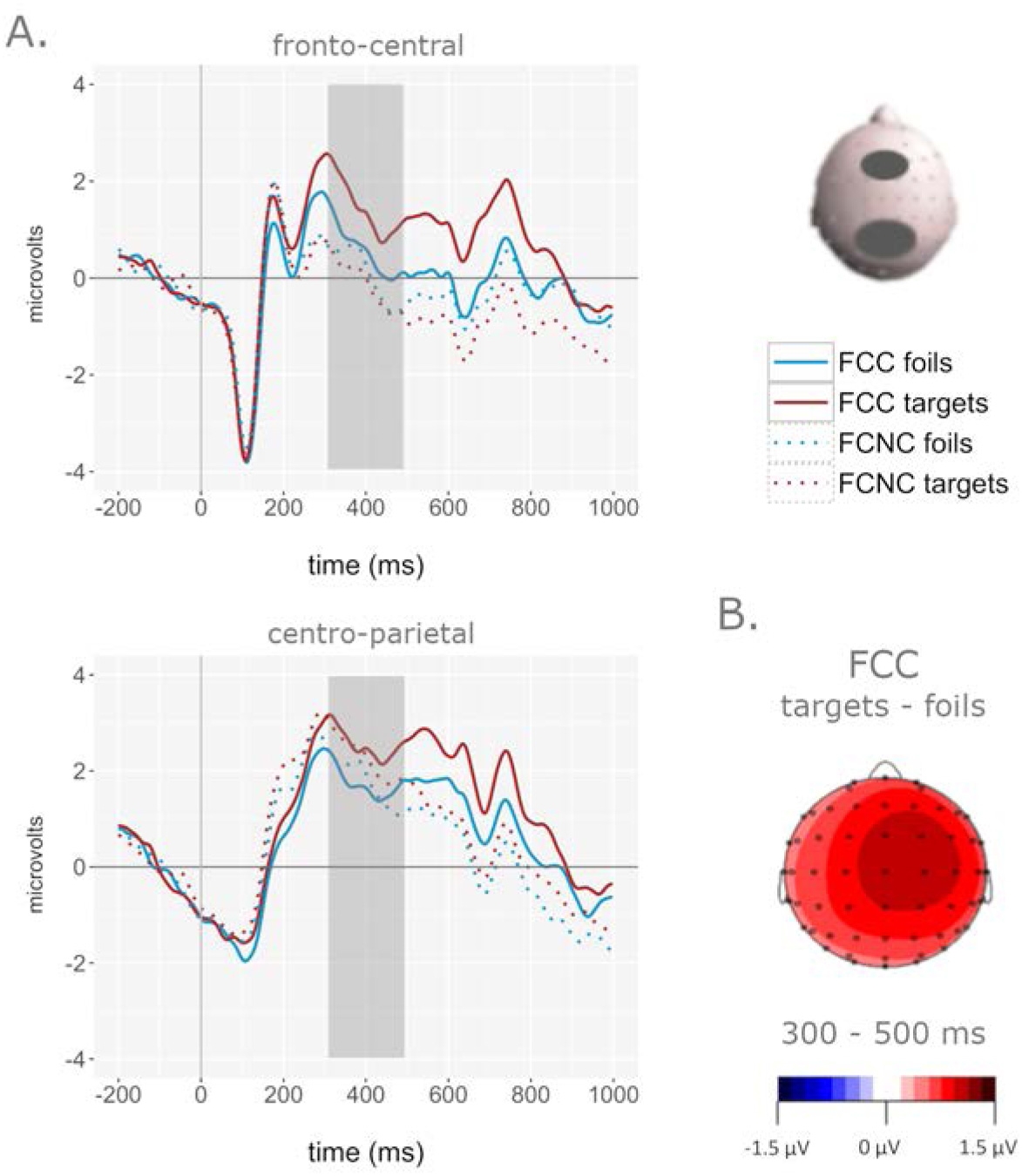
ERP results for the second presentation cycle. A. ERP waveforms at the fronto-central and the centro-parietal electrode cluster for all four conditions. Shaded areas indicate the early analysis time window. B. Topographic distribution of the target vs. foil difference in the FCC condition for the early time window.

**Figure 4.**
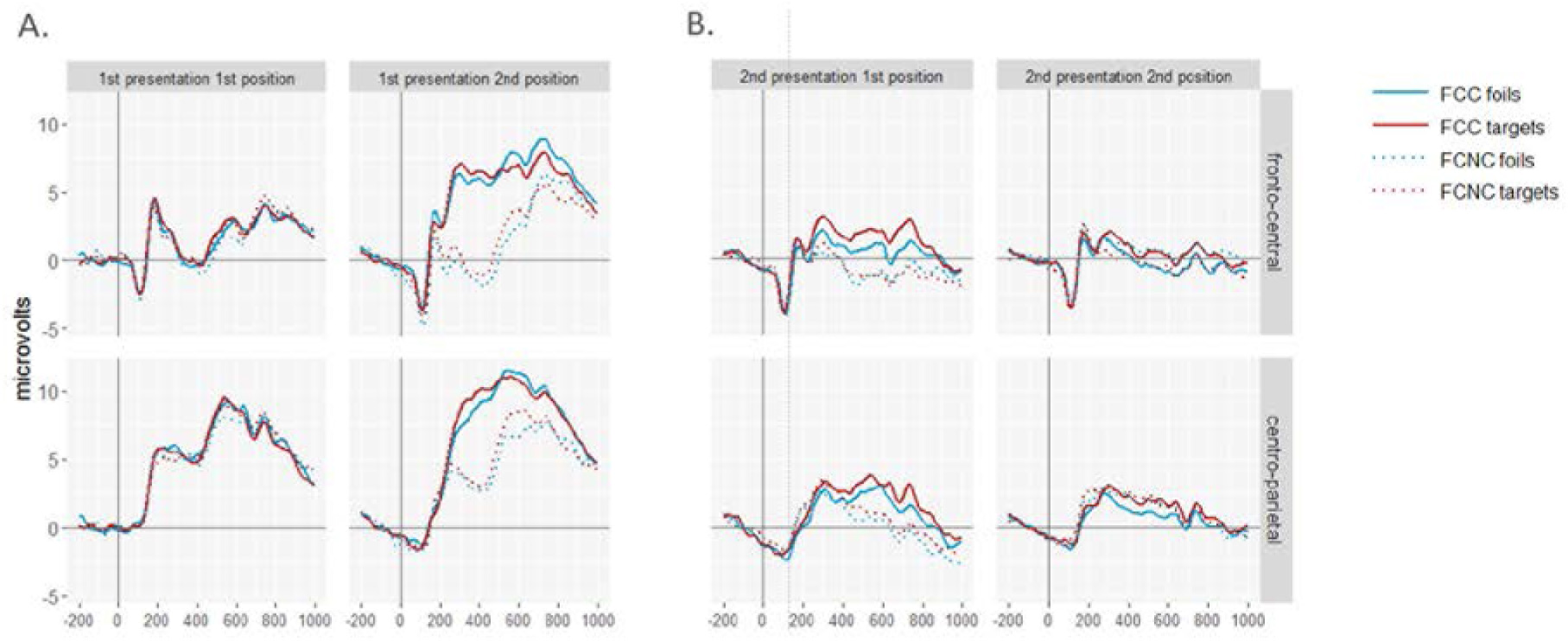
ERP waveforms at fronto-central (upper panel) and centro-parietal (lower panel) electrode clusters for targets and foils in both conditions. A. First presentation cycle, separately for each picture position. B. Second presentation cycle, separately for each picture position.

### Time Window 300-500 ms

A 2 x 2 x 2 ANOVA with the factors Condition, Item Type, and Picture Position was run on mean amplitudes measured over the fronto-central electrode cluster during the second presentation cycle yielding a significant main effect of Condition (F(1,31) = 14.45, p = .001, MSE = 4.69, 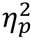 = .32), a significant main effect of Item Type (F(1,31) = 6.22, p = .018, MSE = 1.12, 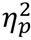 = .17), a significant Condition x Picture Position interaction (F(1,31) = 16.74, p < .001, MSE = 3.03, 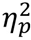 = .35), a marginally significant Item Type x Picture Position interaction (F(1,31) = 4.09, p = .052, MSE = 2.35, 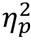 = .12) and as predicted a significant Condition x Item Type interaction (F(1,31) = 5.86, p = .022, MSE = 3.27, 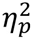 = .16). The latter interactions were not qualified by a Condition x Item Type x Picture Position interaction (F(1,31) = 1.27, p = .268, MSE = 2.63, 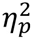 = .04). The main effect of Picture Position (F(1,31) = 0.002, p = .967, MSE = 2.83, 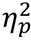 < .001) was also not significant.

To follow-up the significant Condition x Item Type interaction, amplitudes at fronto-central sites to target and foils were compared for each condition separately, collapsed across both picture positions. As planned comparisons revealed, in the FCC condition, targets elicited significantly more positive-going waveforms than foils (t(31) = 3.24, p = .001, d_av_ = 0.27, one-tailed) while the difference was reversed, but not significant in the FCNC condition (t(31) = 0.86, p = .398, d_av_ = 0.08). To follow-up the significant Condition x Picture Position interaction, we compared post-hoc the two conditions for each picture position separately, collapsed across item types. For the first position, mean amplitudes in the FCC condition were significantly larger than in the FCNC condition (t(31) = 5.78, p < .001, d_av_ = 0.66). In contrast, there was no significant difference between conditions for the second picture position (t(31) = 0.39, p = .702, d_av_ = .04). To sum up, in the early time window in the fronto-central ROI, analyses of the second cycle revealed an old/new effect in the FCC condition, but not in the FCNC condition. Moreover, across both item types, there was a significant condition difference in the first but not the second position.

### Time Window 500-650 ms

A 2 x 2 x 2 ANOVA with the factors Condition, Item Type, and Picture Position on the centro-parietal electrode cluster for the second cycle revealed significant main effects of Condition (F(1,31) = 16.25, p < .001, MSE = 4.00, 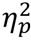 = .34) and Item Type (F(1,31) = 5.28, p = .028, MSE = 3.47, 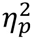 = .15) as well as a significant interaction of Condition x Picture Position (F(1,31) = 36.25, p < .001, MSE = 2.80, 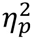 = .54). However, all other interaction effects were not significant (ps ≥ .200).

Due to the significant Condition x Picture Position interaction, we post-hoc compared conditions for each picture position separately, collapsed across item types. For the first position, waveforms in the FCC condition were significantly more positive-going than in the FCNC condition (t(31) = 6.64, p < .001, d_av_ = .77) whereas there was no difference for the second position (t(31) = 0.81, p = .423, d_av_ = .10). To sum up, in the late time window a reliable old/new effect was evident across both conditions. Moreover, a condition difference was observed for the first but not for the second position.

### Multi-level Logistic Regression Model

The final models using the mid-frontal old/new effect as predictor are summarized in Table 1. First, we analyzed whether the single trial target-foil difference score predicts the accuracy of a response separately for both conditions. In the FCC condition, we compared the random intercepts / random slopes model with the random intercepts only model and found that it was not significantly better in predicting response accuracy (*χ*^2^(2) = 0.12, p = .942). Thus, we kept the random intercepts only model, in which the individual predictor difference score was significant (z = 1.87, p = .031, one-tailed). In the FCNC condition, the random intercepts / random slopes model was not better than the random intercepts only model (*χ*^2^(2) = 1.63, p = .442). In contrast to the FCC condition, the predictor difference score was not significant in the latter model (z = 0.94, p = .345, two-tailed). Thus, the target-foil amplitude difference successfully predicts the accuracy of a response only in the FCC condition, but not in the FCNC condition.

**Table 1:**

|  | FCC | FCNC | full model |
| --- | --- | --- | --- |
| Intercept | 1.72 (0.13) | 1.33 (0.14) | 1.73 (0.13) |
| Random effects | 0.42 | 0.55 | 0.48 |
| Difference score | .009 (.005) | .004 (.004) | 0.009 (0.005) |
| Condition |  |  | -0.42 (0.07) |
| Difference score * condition |  |  | -0.005 (0.006) |

Coefficients (standard error) of the final random intercepts models for the early time window.

Second, we tested whether the target-foil difference score better predicted the accuracy of a response in the FCC than the FCNC condition. For this purpose, the centered difference score, centered values of condition and the interaction term of these variables were entered into the model. The random slopes / random intercepts model did not fit the data better than the random intercepts only model (*χ*^2^(9) = 12.01, p = .213). Thus, we kept the random intercepts only model, however, the interaction term was not significant (z = −0.81, p = .421) suggesting that prediction in the FCC condition was not significantly better than in the FCNC condition.

## Discussion

This study set out to provide the (to our knowledge) first neurocognitive evidence in healthy subjects for one core prediction of the complementary learning systems framework (Norman & O’Reilly, 2003), namely that familiarity is more diagnostic in forced-choice corresponding (FCC) tests, in which the familiarity strength of two items can be directly compared, than in other test formats where no direct comparison is possible. As predicted, we showed that the mid-frontal old/new effect, the putative electrophysiological correlate of familiarity-based recognition memory (Rugg & Curran, 2007) is larger in a FCC test format where targets and similar foils are presented in the same trial than in a forced-choice non-corresponding test format (FCNC) where targets are presented together with foils which are similar to other targets from the study phase. According to the CLS framework, the medial temporal lobe cortex (MLTC) generates familiarity signals by assigning highly overlapping representations to similar inputs resulting in small differences in familiarity strength between studied targets and similar foils. As FCC tests allow direct comparison of familiarity strength values of similar items, this test format allows adequate recognition judgments even when only small differences in familiarity strength are accessible. In contrast, in FCNC tests, similar to standard YN formats, familiarity strength values must be compared to a global decision criterion which is problematic when familiarity distributions strongly overlap.

Our study is in line with other studies that show an increase in the accuracy of familiarity-based judgments for FCC tests (Bastin & van der Linden, 2003; Holdstock et al., 2002; Migo et al., 2009). Notably, in the only other study that investigated test format effects in healthy young participants we are aware of (Migo et al., 2009), participants were instructed to exclusively rely on familiarity. This procedure is problematic as it poses high demands on the subjects’ insights into the nature of familiarity and recollection and on their ability to suppress recollection. Thus, our study is the only study which dissociated the FCC and FCNC test formats without relying on subjects’ meta-memory abilities. Moreover, as other studies with patients have revealed contrary results (Bayley et al., 2008; Jeneson et al., 2010), evidence from healthy participants is especially important.

Applying a logistic regression model, we also showed that the target-foil difference wave at fronto-central electrodes in the 300-500 ms time window in a single trial is related to the accuracy of the response in this trial. However, although the single-trial analyses revealed that this was only the case for the FCC condition, i.e. when familiarity is supposed to be highly useful for recognition decisions, it remains unanswered how specific the mid-frontal old/new effect’s predictive value is for the FCC condition as the condition by difference score interaction was not significant.

From a methodological perspective, analyzing old/new difference scores on a single-trial basis opens plenty of possibilities for new research questions. Especially, this single-trial perspective can be informative about the importance of a given neural signature for the outcome of a trial such as the nature or speed of the response over and above the general presence of this signature across all subjects (see Ratcliff et al., 2016, for a similar argument). The study by Ratcliff et al. used an approach similar to multivariate pattern analysis to fit single trial EEG data and found that only late parietal but not early frontal EEG activity is predictive for recognition memory decisions. At first glance, this seems to be at odds with the current results. However, we rather suggest that the use of familiarity and recollection depends on the actual test situation. Here, we created conditions in which familiarity was especially diagnostic and thus was useful in the recognition decision. Accordingly, using data from a source memory task, Noh et al. (2018) extracted an EEG classifier with a spatio-temporal distribution reminiscent of the mid-frontal old/new effect that best distinguished between hits without source judgments and correct rejections implicating that this component reflects a diagnostic familiarity signal.

As old/new differences at parietal electrodes from 500 ms onwards are normally associated with recollection, we also analyzed the time window from 500 to 650 ms. The old/new difference in this time window was not moderated by condition and displayed a similar topographical distribution as in the earlier time window. Prolonged frontal old/new effects that extend beyond 500 ms are not unusual and have been reported in a variety of studies before (e.g., Mecklinger, Brunnemann, & Kipp, 2010; Schloerscheidt & Rugg, 2004; Tsivilis, Otten, & Rugg, 2001). It is rather worth noting that the typical late parietal portion of the old/new effects was not observed in this paradigm. However, interpreting this as a complete lack of recollective processing would certainly be exaggerating given the relatively high performance levels in both conditions. However, it is also conceivable that recollection played a minor role as the use of long study lists may have reduced its contribution (Cary & Reder, 2003). Another explanation for the absence of the late parietal old/new effect in the FCNC condition is that recall-to-reject processing (Rotello, Macmillan, & Van Tassel, 2000), i.e. recall of item details of the originally studied picture upon the presentation of foils, might have alleviated differences between targets and foils, thereby disguising the late parietal effect. Supporting this interpretation, Migo et al. (2009) found evidence for reliance on recall-to-reject in a remember/know variant of their experiment in which participants were asked to verbalize their decision process. A third possibility is that recollective processing was spread across all four pictures of the trial sequence and the intervals between the pictures. Consequentially, recollection was presumably less time-locked to stimulus onset and therefore not observable in the ERPs. Note that temporal smearing is less likely for the mid-frontal old/new effect as familiarity is assumed to be elicited fast, more automatically, and with less temporal jitter.

Repeating pictures within a trial in the test phase seems to be both, a methodological strength and caveat of this study. As outlined in the Introduction, the necessity of a sequential presentation in order to obtain separate ERPs for targets and foils implicated that the difference between the two conditions has an effect only after each picture has been presented at least once. In support of this view, it can be seen in Fig. 4A that there was indeed no condition difference during the first picture position of the first cycle. Thus, a second presentation cycle was inevitable for a sound comparison between the two conditions. However, a within-trial repetition also bears the risk that differential processing in the first cycle affects processing in the second cycle and ERPs have to be interpreted carefully. As can be seen in Fig 4A, repetition of similar pictures in the FCC condition leads to a positive shift in the waveforms during the second picture in the first cycle (and also persisting to the first picture of the second cycle) presumably due to repetition priming (Penney, Maess, Busch, Derrfuss, & Mecklinger, 2003; Penney, Mecklinger, & Nessler, 2001). Consistent with this repetition priming view, this ERP difference between first and second presentation in the first cycle is virtually absent for the dissimilar pictures in the FCNC condition. Importantly, a putative priming mechanism in the FCC condition cannot explain larger target-foil differences in this condition as it should affect the processing of target and foils to the same extent. Thus, we feel save to conclude that the larger mid-frontal familiarity effect in the FCC condition is not an artefact of differential processing during the first cycle.

Our results also have implications for discussions on the functional significance of the mid-frontal old/new effect. Mirroring the imprecision in the definition of the familiarity process itself (e.g., feeling of “knowing” or recognition without recollection of details), the exact functional significance of the mid-frontal old/new effect remains elusive. One suggestion was that the mid-frontal old/new effect merely reflects differences in conceptual fluency between studied and non-studied items (Paller et al., 2007). However, our results add to other findings (Bader & Mecklinger, 2017; Bridger, Bader, Kriukova, Unger, & Mecklinger, 2012) strongly speaking against this explanation. We observed the mid-frontal old/new effect only in the FCC but not in the FCNC condition although across all items in a test list, differences in conceptual fluency between targets and foils were equated between conditions as foils in the FCNC condition were also similar to another picture from the study list. More precisely, as we assume that differences between the familiarity distributions of targets and foils are of the same size in the two test display conditions, the current results suggest that the mid-frontal old/new effect reflects a task-adequate and fast *assessment* of the familiarity signal, not the signal associated with the pure familiarity strength value itself. This is in line with the assumption that familiarity is multiply determined. In previous studies (Bader, Mecklinger, Hoppstädter, & Meyer, 2010; Bridger, Bader, & Mecklinger, 2014; Wiegand, Bader, & Mecklinger, 2010), we showed that early ERP old/new effects with parietal maxima are likely associated with *absolute* familiarity which signals the strength of the memory representation at a given time point. In contrast, more frontally distributed old/new effects, as revealed in the FCC condition in the current experiment, were linked to the *relative* increment of the familiarity strength value for an item due to a specific recent encounter. Thus, relative familiarity is not an integral characteristic of a stimulus but the by-product of an assessment process. This is also supported by findings that the mid-frontal old/new effect is specifically tied to explicit recognition memory tasks, in which a discrimination between old and new items is task-relevant, i.e. if familiarity strength has to be assessed to guide recognition judgments (Ecker & Zimmer, 2009; Guillaume & Tiberghien, 2013; Küper et al., 2012). The mid-frontal old/new effect is usually not present in tasks requiring non-mnemonic judgments as for example judgments of lifetime exposures (Yang et al., 2019) or during heuristic decision making (Rosburg et al., 2011). In these tasks, the mid-frontal familiarity effect is replaced by a more posterior effect, resembling the N400, an ERP index of semantic processing.

In this context, the question also arises whether the test display affects processing in the MTLC as suggested by the CLS (Norman & O’Reilly, 2003) or by other brain structures involved in familiarity judgements. Indeed, as we have discussed previously (Bader & Mecklinger, 2017), it seems more likely that the comparison of familiarity strength values, i.e. the assessment of the relative increment in familiarity and the requirement to make explicit recognition judgements, is mediated by the prefrontal cortex (PFC) while the MTLC might be more involved in the generation of the familiarity signal itself, i.e. absolute familiarity. Thus, even though inferences from scalp distributions of ERP effects on underlying brain systems have to be made with caution, we think that differences in the mid-frontal old/new effect between FCC and FCNC displays most likely are due to differences in prefrontal activity related to processes responsible for familiarity-based episodic decision making. Such a relationship between ventro-lateral PFC activity and the mid-frontal old/new effect was recently demonstrated by an EEG-informed fMRI study (Hoppstädter, Baeuchl, Diener, Flor, & Meyer, 2015).

In conclusion, showing that the usefulness of a familiarity signal in a recognition memory task depends on the test format, we provide evidence in favor of the CLS model for recognition memory. Moreover, the current results suggest that the mid-frontal old/new effect does not reflect the mean difference in absolute familiarity strength between old and new items but reflects the assessment of the familiarity signal to pursue episodic recognition memory judgments.

Data is available on: https://osf.io/n2w83/?view_only=d28941463fce4420a47ac6cad3804acd

## Acknowledgements

The authors thank Alexander Hauck, Kristin Pfaff, and Lisa Riedel for assistance with data collection and analyses.

